# Frontal-midline theta and posterior alpha oscillations index early processing of spatial representations during active navigation

**DOI:** 10.1101/2023.04.22.537940

**Authors:** Yu Karen Du, Mingli Liang, Andrew S. McAvan, Robert C. Wilson, Arne D. Ekstrom

## Abstract

Previous research has demonstrated that humans combine multiple sources of spatial information such as self-motion and landmark cues, while navigating through an environment. However, it is unclear whether this involves comparing multiple representations obtained from different sources during navigation (parallel hypothesis) or building a representation first based on self-motion cues and then combining with landmarks later (serial hypothesis). We tested these two hypotheses (parallel vs. serial) in an active navigation task using wireless mobile scalp EEG recordings. Participants walked through an immersive virtual hallway with or without conflicts between self-motion and landmarks (i.e., intersections) and pointed toward the starting position of the hallway. We employed the oscillatory signals recorded during mobile wireless scalp EEG as means of identifying when participant representations based on self-motion vs. landmark cues might have first emerged. We found that path segments, including intersections present early during navigation, were more strongly associated with later pointing error, regardless of when they appeared during encoding. We also found that there was sufficient information contained within the frontal-midline theta and posterior alpha oscillatory signals in the earliest segments of navigation involving intersections to decode condition (i.e., conflicting vs. not conflicting). Together, these findings suggest that intersections play a pivotal role in the early development of spatial representations, suggesting that memory representations for the geometry of walked paths likely develop early during navigation, in support of the parallel hypothesis.

## 1. Introduction

Spatial navigation is an essential ability to survival (e.g., finding food or shelter). In any environment, forming an accurate spatial memory will guide the navigator to make effective decisions and plan for the future steps. In such a dynamic process to develop spatial memories, various navigational cues are involved: 1) external landmarks for orientation (termed “allothetic cues”), and 2) optic flow and other body-based cues for keeping track of distance (termed “idiothetic” cues). Differential weighting of allothetic and idiothetic cues is commonly observed among many tasks involving cue combination and competition (Du, Mahdi, et al., 2016; Du, Spetch, et al., 2016; Harootonian et al., 2020; Mou & Zhang, 2014; L. Wang et al., 2018), which may suggest evidence for an accumulation process for selecting and assigning attributes to spatial memories. For example, a building with unique colors that suggest value as a landmark may be weighted more by the navigator than other more drab buildings on that street so that it may act as an anchor for memory and orientation (Janzen & Van Turennout, 2004; Walter et al., 2022).

As part of combining idiothetic (self-motion cues like body movements and optic flow) and allothetic cues (proximal and distal landmarks), humans’ spatial memory may not accurately reflect their physically experienced reality (Du et al., 2020; Harootonian et al., 2020; McNamara & Diwadkar, 1997; Mou & McNamara, 2002; Shelton & McNamara, 2001; Tversky, 1981; Warren et al., 2017). In everyday life, if we expect a landmark at a certain location and discover it is at a different location, we adjust our internal estimate of our current location and bearing based on reorienting to the visual landmark. In certain situations, memory distortions may violate physical Euclidean geometry or even topology (Du et al., 2023; Ericson & Warren, 2020; Warren et al., 2017). In this way, combinations of idiothetic and allothetic cues may occur in a way that results in distortions of physical space, particularly if landmarks conflict with the expected position based on idiothetic cues. The combination of allothetic and idiothetic cues is difficult when studying regular environments (those that occur in everyday life) and therefore it is necessary to turn to immersive virtual reality to create situations in which idiothetic and allothetic cues can be mismatched.

In a recent study (Du et al., 2023), participants walked through hallways in immersive virtual reality and pointed back to the start of the hallway. Unbeknownst to the participants, some conditions involved removing an expected visual intersection on a crossed path or adding a “false” intersection on an uncrossed path. The results demonstrated large memory errors regarding the shape of the path walked by participants when experiencing either the absence of an expected intersection or the presence of unexpected intersections. Specifically, when the intersection between two path segments was visually hidden in a crossed path, participants pointed in a manner suggesting they had experienced an uncrossed path. In contrast, when participants were shown non-functional, false intersections on an uncrossed walked path, participants pointed in manner suggesting a crossed path. Such findings suggest that when combining conflicting idiothetic and allothetic cues, memory distortions may arise that violate the Euclidean properties of physical space. In this previous study, participants indicated their memory for paths by pointing to where they thought they started, which would be subject to the distortions imposed by our experimental manipulations. By asking participants to point at the end of the path, however, we gained only incomplete knowledge of the participant’s assumptions and ideas about their knowledge about the path. If we had asked participants to point (or even draw) the path throughout navigation, this would have both disrupted their natural navigation-related movement and also could have influenced their subsequent retrieval (known as test-retest effects; also see L. Zhang & Mou, 2017). Therefore, to provide continuous insight into participants’ build-up of knowledge about paths, we recorded simultaneous scalp EEG, which we used to test rival hypotheses about when participant hypothesized representations for paths first emerged.

Although previous research has revealed that humans’ spatial memory may be altered from the real environment and with prominent errors, there are competing ideas about how distortions in spatial memory may arise. One idea is that navigation involves a gradual accumulation of idiothetic cues for distance and direction and eventual integration with allothetic landmark cues (O’keefe & Nadel, 1978; Siegel & White, 1975). For example, O’Keefe and Nadel (1978) described the process of “exploration” as fundamental to establishing direction and distance, with allothetic landmarks providing for matches or mismatches once the environment is sufficiently learned. Once the mismatches can be resolved, a cognitive map can be established. Similarly, Siegel and White (1975) discussed the idea of route information, which involves turns and distances, as coming before integration with landmarks and a schematic “map” can be established. These, and later models, postulate that self-movement (idiothetic) cues provide information about direction and distance for the formation of the cognitive map with landmarks providing a means of correction once the map is established through sufficient exploration (McNaughton et al., 2006; Redish & Ekstrom, 2013). Such models are broadly consistent with the idea that initially, path integration information may be fairly accurate but accumulates error over time. As such, landmarks play an important role in correcting or resetting the path integration systems (Fujita et al., 1993; Harootonian et al., 2020; Loomis et al., 1999; R. F. Wang, 2016).

Another possibility is that much of learning about spatial layouts occurs in parallel and involves an active process of comparing available information with current predictions (Harootonian et al., 2020; Ishikawa & Montello, 2006; Sjolund et al., 2018; H. Zhang et al., 2014). One class of these models suggests that idiothetic and allothetic information may be acquired early and in parallel, allowing for active integration and comparison (Ishikawa & Montello, 2006; H. Zhang et al., 2014). Another class of models, which rely on a Bayesian probabilistic mathematical framework, suggest that participants integrate idiothetic (self-motion) and allothetic (landmark) information in an “optimal” fashion by combining unreliable cues (noisy self-motion cues and unstable landmarks) to reduce the variance in estimates of position (Chen et al., 2017; Nardini et al., 2008; Newman et al., 2023; Sjolund et al., 2018; Zhao & Warren, 2015). Some such Bayesian models additionally incorporate participants’ knowledge obtained from previous trials, which may include guesses about distance and direction that can be (either correctly or incorrectly) applied to the current trial (Harootonian et al., 2022).

According to such models, participants should incorporate idiothetic and allothetic cues early during learning to make predictions about the shapes of paths that they are walking. Additionally, participants may even employ information from past trials to inspire their internal estimates about the path shapes of current trials. Together, these can lead to multiple predictions and representations that the participant generates in an attempt to reconcile guesses and estimates of the current path shape in the presence of conflicting information.

Here, we sought to test the competing predictions of the broad class of “serial” vs. “parallel” navigation learning models. As summarized above, serial navigation models postulate the primacy of idiothetic cues to navigation, followed by matching or mismatching allothetic cues, leading to a single spatial representation. Parallel models argue that idiothetic and allothetic information are often compared and integrated early and in parallel, along with “guesses” about path shape from past trials, leading to potentially multiple representations that are ruled in or out. Simultaneous wireless scalp EEG offers a direct opportunity to tacitly observe the build-up of participants’ putative representations about paths without having to explicitly ask them behaviorally about such knowledge, which could have influenced their subsequent retrieval. We employed the simultaneously oscillatory signal present with both frontal-midline theta and posterior alpha to determine: 1) how participants incorporated landmarks (in this case, intersections) into their current representations and 2) when during navigation (early vs. late) neural signals showed evidence of distinguishing between guesses about path shape. Both of these signals provide indices into participants memories and cognitive processes relevant to understanding how participants might be representing path shape based on incoming landmark and idiothetic cues. We discuss previous work on frontal-midline theta and posterior alpha oscillations in more detail shortly.

Here, we used a task from the aforementioned “hallway” study (Du et al., 2023). Specifically, we asked participants to physically navigate in an immersive virtual hallway with a head-mounted display (HMD) and point to the start location of the hallway when they reached the end (Figure 1A, B, C). The eight different hallways were designed to include either one true intersection in a crossed shape (TI-C, “true intersection – cross” condition) or to include two false intersections in a non-crossed shape (FI-NC, “false intersection – no cross” condition). We included a period of active walking of four legs of a path, including passing visible intersections, and a period of imagination in which participants were instructed to imagine their start location directly before they pointed to the start position. Scalp EEG was recorded during the walkthrough of the hallways. Through the lens of time-frequency analysis, competing hypotheses can be evaluated by comparing oscillatory patterns at critical intersections between conditions. We focused on oscillations in theta band (4-8 Hz) in frontal-midline regions and alpha band (9-12 Hz) in posterior regions based on previous work using wireless scalp EEG showing that frontal-midline theta and posterior alpha are two sources of salient oscillations during navigation (Liang et al., 2018). The results from this paper demonstrated a dissociation between frontal-midline theta during movement and posterior alpha during imagination, and coupled with another paper, which showed posterior alpha oscillations and frontal-midline theta oscillations relevant to distance coding using wireless scalp EEG (Liang et al. 2021), form part of the foundation for the current work. Previous research has also shown that frontal-midline theta oscillations relate to the encoding and retrieval of spatial information (Chrastil et al., 2022; Liang et al., 2018, 2021), with posterior oscillations in parietal and occipital regions relating to increased attention prior to decision making (Benedek et al., 2014; Chrastil et al., 2022; Klimesch et al., 1998; Sauseng et al., 2005), suggesting that theta and alpha oscillations are involved in the latent components supporting successful navigation. Together, these findings demonstrate their relevance to navigation, memory, and decision-making suggesting their potential usefulness as a proxy into what sorts of information participants might be using to build up representations about different paths as they navigated. Therefore, we focused our analyses on frontal-midline theta during walking and posterior alpha during imagination prior to pointing.

**Figure 1.**
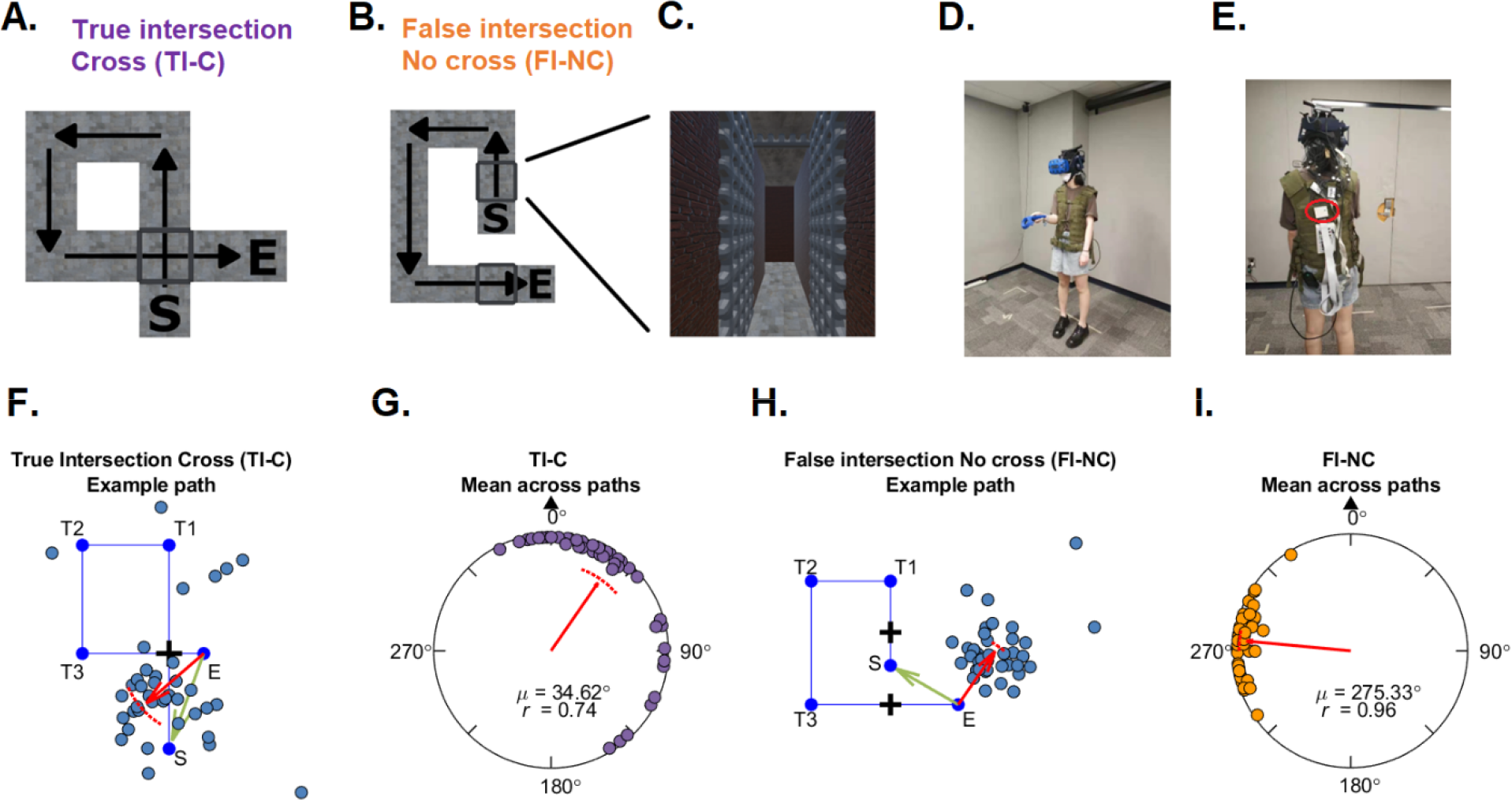
(A-B) Examples of hallways for True intersection – Cross (TI-C) and False intersection – No cross (FI-NC) conditions. C shows a leg of the hallway with the obscuring gates at a false intersection from a participant’s perspective. (D, E) Experimental testing space, HMD, and EEG recording devices. The circled device is the MOVE module. (F, H) Behavioral pointing responses in example paths. Each dot indicates the pointing direction of one participant across trials for the path. The red arrow indicates the circular mean direction of the pointing directions across all participants. The arc above the mean direction indicates the 95% confidence interval of the mean direction. Green arrows indicate the correct pointing direction according to Euclidean geometry. S: Start. T1-3: Turn 1-3. E: End. (G, I) Mean of circular pointing errors (μ).

In the current study, we used P-episode to quantify the presence of theta and alpha oscillations. The methods segment neural timeseries into non-oscillatory and oscillatory portions and use the percentage of oscillatory portions as a proxy for the prevalence of neural oscillations. According to the models discussed previously, EEG data were analyzed to contrast two competing predictions. According to the serial hypothesis, idiothetic information is primary and accumulates prior to landmark information, then oscillatory prevalence would distinguish between true and false intersection conditions later in learning, possibly during the imagination phase. This is because participants should rely primarily on idiothetic cues to establish the direction and distance of the start position, possibly combining mismatching landmarks later during the path. In contrast, parallel models would argue that idiothetic and allothetic integration occur early and continuously during learning as part of a Bayesian processes. Therefore, serial models would appear to predict that neural oscillations would discriminate between conditions later in learning, likely during the imagination period when participant attempt to reconcile idiothetic and allothetic information, while parallel models predict that neural oscillations would discriminate between conditions early in trials, particularly at intersections.

We tested parallel vs. serial models of navigation using neural representational similarity and classification approaches. If neural representations reflecting spatial knowledge occur late during navigation, we would expect that condition-specific neural representations would emerge near the end of a trial when participants remembered details about the path, which in this case would involve the imagination phase. This is because the imagination phase was the final period before participants pointed to their start position and would be a moment in which participants would be required to remember, as best as they could, each of the four legs and two intersections of the hallway to compute their global path. In contrast, if neural representations reflecting spatial knowledge emerge early during navigation based on evidence accumulation related to decision making and memory processes that occur continuously during navigation, we would expect that representational similarity analyses during earlier walking phases (e.g., comparing oscillatory activity during the first and second intersection and how this related to subsequent pointing) and classification analyses might reveal significant information about condition and even specific paths.

## 2. Methods

### 2.1. Participants

Forty-eight participants (26 female and 22 male) were recruited from the University of Arizona undergraduate Psychology program and surrounding Tucson city area whose age ranged from 18 to 36 years with a mean age of 22.96. One participant (female) withdrew from the experiment in the middle due to cybersickness. Four of the participants (three female and one male) did not complete all the trials due to technical issues of the wireless virtual reality (e.g., HMD disconnection). We obtained complete behavioral data from the remaining 43 participants. Due to technical issue of the mobile EEG system or the experimenter error, the EEG recordings of four participants (one female and three male) were incomplete. We obtained complete EEG recordings from 39 participants (21 female and 18 male).

All participants were compensated at a rate of $20 per hour. All participants had normal or corrected-to-normal vision, normal or corrected-to-normal hearing, and reported no history of cardio-vascular problems or motion sickness. Written consent was obtained before the experiment, and the methods were approved by the Institutional Review Board (IRB) at the University of Arizona.

We report how we determined our sample size, all data exclusions, all inclusion/exclusion criteria, whether inclusion/exclusion criteria were established prior to data analysis, all manipulations, and all measures in the study. We based our sample size (∼ 20 for each test order, see the following sections) on previous research which examined pointing responses in similar behavioral tasks (Du et al., 2023) and mobile scalp EEG involving spatial information processing (Liang et al., 2021).

### 2.2. Apparatuses

The experiment was conducted in a physical room approximately 6 m × 6 m in size with the navigable virtual environment being approximately 5 m × 5 m in size. The virtual environment and experimental tasks were built in Unity 3D engine (Unity Technologies ApS, San Francisco, CA) using the Landmarks virtual reality navigation package (Starrett et al., 2021). Participants experienced the fully immersive virtual environments via the use of an HTC Vive Pro head-mounted display (HMD) fitted with an HTC Wireless Adapter (HTC, New Taipei City, Taiwan) that allowed for untethered, free ambulation. The Vive Pro displayed stimuli at a resolution of 1140 × 1600 pixels per eye with a 110° field of view (FOV) refreshed at a rate of 90 Hz. The Wireless Adapter allowed for participants’ orientation and position to be tracked for up to 7 m, with data being delivered over a 60 GHz radio frequency. A handheld HTC Vive controller (HTC, New Taipei City, Taiwan) was used to record responses from participants and interact with the virtual environment. The tasks were run on a custom-built computer with an NVIDIA GeForce Titan Xp graphics card (NVIDIA Corp., Santa Clara, CA).

The continuous EEG recordings were acquired with a 64-channel BrainVision ActiCAP system, which included a wireless transmission MOVE module and two BrainAmp amplifiers (BrainVision LLC). We recorded from 64 active electrodes, placed on the scalp according to the International 10–20 system. The reference electrode was located at FCz; no online filter was applied to the recordings. Before the experimenter started the recordings, the impedances of all 64 electrodes in most participants were confirmed to be below 60 kΩ. For 14 electrodes across all participants (0.56%; in four participants), impedances were above this criterion (range: 61-207 kΩ), with 6 of these channels interpolated from other channels (see Section 2.5.1). The sampling rate was set to 500 Hz. The MOVE module and the control box, as well as the battery pack that powered the HMD, were clipped onto the back or shoulder areas of a vest which the participant wore for the whole experimental session (Figure 1D, E).

Across paths. The black triangle indicates the pointing direction consistent with Euclidean geometry (0°). *R* is the mean resultant length of all pointing angles.

### 2.3. Materials and design

Participants received a total of 96 trials with eight path shapes in two conditions (visibly intersecting or not; see Figure 1A, B, C). Each condition was presented in two blocks of 24 trials in an alternating order. The condition order (i.e., which condition was first tested) was counterbalanced across participants. For each condition, there were four different path shapes (see Du et al., 2023, Table S1 and Figure S1 for details). Each path had four segments (“legs”) with three clearly visible 90° turns (left/right) requiring rotating the body 90°. The total path lengths, pointing angle from the end location to the start location, and Euclidean end-to-start distances were matched for each pair of the paths. For example, Path 1 was the same as Path 5 in terms of the path length and end-to-start distance, and the left-turn version of Path 1 had the same correct pointing angle as the right-turn version of Path 5. Each path shape was presented for 6 trials in each block. Within each block, the trials representing a specific condition were presented in a random order such that the paths were completely randomized. We randomized the paths within condition blocks so that participants would be unable to guess the global path as they navigated. Trials with the same path shape were never presented twice in a row.

There were eight different hallway models (i.e., 4 path shapes × 2 left/right turning directions) in the experimental program for each condition. In each block, each of these eight hallway models was presented three times with different orientations in the virtual environment. For each participant, the three presentation orientations of each hallway model were randomly chosen from four orientations with respect to the physical room (i.e., 0°, 90°, 180°, or 270° rotated clockwise from the model’s original orientation). For these hallway models, different textures and colors were applied to the walls to ensure that each hall was distinct and provided a sense of optic flow to participants.

For the visibly intersecting hallways, a crossing between the first and the last segments of the path was presented to the participants during walking, with a grated gate to each direction of the intersection (Figure 1C). The gates were shown in an “open” position along the current walking direction and a “closed” position for the orthogonal directions. In this condition (“true intersection - cross”, TI-C), the participant experienced the intersection as they would in the real world. For the non-intersecting hallways, two false intersections were shown in the first and the fourth legs of the hallway (“false intersection - no cross”, FI-NC condition). They served no function (other than experimental manipulation) as they were not connected to any other parts of the hallway. The two conditions were comparable because in both situations, participants went through the grated gates twice in two orthogonal directions. Participants did not receive any specific verbal instructions for recognizing the intersections (e.g., “this is an intersection”) because these were visually recognizable and salient features of the hallways. For a demonstration of what a scene looked like to a participant, please see our example videos at https://osf.io/8jhav.

Two additional path shapes were used in the practice prior to the formal trials to familiarize the participants with the paradigm. Both had two paths of equal distance (2.55 meters), with one containing a single 90° left turn and the other containing a single 90° right turn. The paths were presented by showing hallways with a wall texture different from those in the formal trials. No intersection was presented in these practice hallways.

### 2.4. Procedure

Participants were tested individually. After giving their written consent, the participants first performed four practice trials (which followed the same procedure as the formal trials) with the wireless HMD and the controller. In this practice phase, the experimenter made sure that the participants were familiar with the paradigm and followed the instructions such as looking forward, avoiding stopping, and not leaving the designed path. Then the experimenter placed the scalp EEG recording electrodes on the participant’s head and applied the electrode gel to reduce impedance and assure the quality of recordings. After that, participants were fitted with the HMD again and started the formal trials.

On each trial, participants were instructed to rotate slowly in place while verbally calculating subtractions for approximately 10 seconds (e.g., “Please rotate leftward in place and verbally count backward from 100 by 3”). This procedure was intended to disorient participants from any previously gained spatial knowledge. The participants were then presented with a dark environment and were asked to search for and walk to a red platform with a directional arrow.

When they arrived at the platform, it turned green, and then they were asked to face the same direction as the arrow. The platform indicated the start of the path, and the participants were asked to remember this position while walking. When participants pressed a button on the controller, the platform disappeared, and the hallway appeared. At this point, the participants stood at the start of the hallway. They walked through all four hall segments until they arrived at the end (indicated by an ending wall). Once at the end, the hallway disappeared, and the participants stood still and were asked to imagine the start of the hallway for 5 seconds (“Imagine where the start is. Stand still.”). The length of this phase (5 seconds) was determined by the Unity program and varied slightly between trials (range: 4962-5586 ms, mean length = 4991.50 ms, SD = 7.52 ms). Then the participants were asked to point to the start position of the hallway with a virtual laser pointer (using the controller). The position where the laser pointer touched the floor was recorded, providing a measure of both distance and direction of the remembered start location relative to the end location. The participants were allowed to turn around if they wanted to. They were instructed to point as accurately as possible, and that response time would not be recorded. After making their pointing response, the participants were asked to rate their confidence for their pointing accuracy by adjusting the length of a dark-grey rectangular bar in respect to a separate light-grey bar. The longer the dark-grey bar, the more confidence expressed, and vice versa. The ratio between the dark-grey bar and the light-grey bar was the confidence rating score (ranging from 0 to 1), with the initial value being set to 0.5 (see example videos at https://osf.io/8jhav). To assure the quality of EEG recordings, participants were instructed to minimize extra body movements such as moving their heads or talking during the walking, imagination, and pointing phases. No part of the study procedures was pre-registered prior to the research being conducted.

### 2.5. EEG data analysis

#### 2.5.1. EEG preprocessing

EEG preprocessing and analyses were performed with EEGLAB (Delorme & Makeig, 2004) and customized code in MATLAB. Only one participant’s continuous data underwent re-referencing to average because their reference electrode was malfunctioning. All other participants’ data was referenced to FCz as during data collection. The signals were then band-pass filtered at 1-50 Hz. Before any further preprocessing steps, the data portions of disorientation phase on each trial, the periods of walking to the start platform, inter-trial intervals, and the rest phase between test blocks were removed.

The remaining data (the start of walking phase --- the end of confidence rating phase) structure was kept continuous by concatenating all trials. Then, Independent Component Analysis (ICA) was performed to correct artifacts. We used an automatic component selection procedure, ICLabel (Pion-Tonachini et al., 2019), to avoid experimenter bias in identifying noisy components. Components were rejected automatically if they had labels of “Muscle,” “Eye,” “Heart,” “Line Noise,” or “Channel Noise” and if their probability was higher than 90% for being one of those labels. On average, 6.26 (9.78% of all components, SD = 2.56) components were rejected. After ICA, the kurtosis of each channel was calculated. Channels with kurtosis greater than 3 standard deviations from the mean were rejected and interpolated. In addition, 11 channels in 7 participants were interpolated (although the kurtosis for these channels were all within 3 standard deviations). Together, each participant had 2 to 10 interpolated channels (3.13-15.63% of all channels) with 7.15 interpolated channels (11.18%) on average.

#### 2.5.2. EEG epoching and segmentation

We examined the periods during walking and imagination. For the walking periods, we separated the whole period into four different events: the first intersection area in Leg 1 (starting to see the intersection --- exiting the intersection area before the first turn), Leg 2 (the first turn --- before the second turn), Leg 3 (the second turn --- before the third turn), and the second intersection area in Leg 4 (starting to see the intersection --- exiting the intersection area before reaching the end). For all events, the times of onsets and offsets were identified by the EEG event markers recorded during the experiment. For the first and the second intersection events, we extracted the period [-2000 ms, 0] before the end of the event on each trial. For the Leg 2 and Leg 3 events, we extracted the period [-1000 ms, 0] before the end of the event on each trial. For the imagination phase, we extracted the period [-4000 ms, 0] before the end of the event on each trial. These period lengths were determined by the recorded lengths of each event to include as many usable trials as possible and exclude extra potential noise sources such as body turning or moving arms at the beginning of the event. The mean length of the Intersection 1 period was 3721.18 ms (± 563.09 SD) in a range of 2012-11298 ms. The mean length of the Leg 2 period was 2868.54 ms (± 590.93 SD) in a range of 1188-6800 ms. The mean length of the Leg 3 period was 3714.12 ms (± 671.62 SD) in a range of 1592-10454 ms. The mean length of the Intersection 2 period was 4276.40 ms (± 594.67 SD) in a range of 2290-14354 ms. We only epoched the period that could be identified in the segment with minimal body rotations or extra movements because we observed during the experiment that on many trials, some participants tended to rotate slowly or pause at the turn areas (although they were instructed to try their best not to stop and to keep moving). This may cause body sway, velocity changes and accidental, unexpected EEG signal changes which may be attributed to abrupt body movements. Similarly, at the beginning of the imagination phase, most of the participants moved their arms down and read the texts on the screen. According to our observations, some participants also swayed their bodies a little because they just stopped walking. To exclude extra noise caused by body movements or reading texts, we did not include the first 1 second of the imagination phase but use the last 4 seconds in our analysis.

For the Intersection 1 event, 3 trials were excluded due to HMD disconnection (1 trial) or extreme length (shorter than 2 seconds; 2 trials). For the Leg 2 and 3 events, 2 trials were excluded due to HMD disconnection. For the Intersection 2 event, 2 trials were excluded due to HMD disconnection. No trial was excluded for the imagination event.

#### 2.5.3. Time-frequency analyses and detection of oscillatory activity

We extracted power and P-episode as the dependent variables for the following analyses. Power estimation across time within a certain period indicated the overall activation level of scalp EEG signals while P-episode indicated the scalp oscillatory patterns in this period. We did not conduct the event-related potential (ERP) analysis across trials because the trial lengths varied for each epoched period, making averaging waveforms potentially problematic due to missing data and potentially incompletely aligned epochs.

We estimated the power during each event period with 6-cycle Morlet wavelets using code from Whitten et al. (2011). We sampled frequencies from 1 to 30 Hz: 1-3 Hz for delta band, 4-8 Hz for theta band, 9-12 Hz for alpha band, and 13-30 Hz for beta band. No baseline correction was applied to the power estimates. Mean power for each band was measured as the power averaged across time points within the event period and across frequencies within a band.

Based on the estimated power at each frequency, we used the Better OSCillation method (BOSC, Whitten et al., 2011) to detect oscillatory activity in the signal. For each participant, we used the average of estimated power across all trials as the background estimation. Then we used the 95% confidence level for the power threshold and three cycles as the duration threshold in BOSC to determine whether the signal at each time point during an event epoch was oscillatory (1) or not (0). The resultant proportion of oscillations across all time points, *P-episode*, indicates the prevalence of oscillatory activities during this event (for this method, also see Liang et al., 2018).

We selected 12 frontal channels and 18 posterior channels as our regions of interest (ROIs) as in previous research (Liang et al., 2018). We included AFz, AF3, AF4, F3, F1, F2, F4, FC3, FC4, FC1, and FC2 in the frontal cluster. We included Pz, P3, P7, O1, Oz, O2, P4, P8, P1, P5, PO7, PO3, POz, PO4, PO8, P6, P2, and Iz in the posterior cluster. For the power estimates and P-episode, we calculated the mean across electrodes within each regional cluster, respectively. As planned comparisons, we used paired t tests to compare the mean power and P-episode between TI-C and FI-NC conditions.

#### 2.5.4. Representational similarity analysis between events

To examine the relationships between events within each condition (TI-C or FI-NC), we performed a representational similarity analysis using the P-episode (which indicated the oscillatory activities) between events within our ROIs. We used P-episode instead of power in this analysis because P-episode can better account for the background spectrum, which could vary between conditions and produce inflated correlations. We specifically compared (a) the first and the second times when participants walked through the intersections (b) the periods when they walked through hallway segments without any intersections. We correlated the P-episode matrices (number of electrode × 1-30 Hz frequency in each ROI) (a) between the events Intersection 1 and Intersection 2 and (b) between the events Leg 2 and Leg 3 (by Pearson correlation) within each trial. Then the correlation coefficient was transformed to Fisher’s Z scores as Z = ln[(1 + r) / (1 - r)] / 2. Similar methods have been used with magnetoencephalography (MEG) recordings and intracranial EEG (Manning et al., 2012; Wardle et al., 2016).

#### 2.5.5. Representational similarity analysis between P-episode and pointing error

To further examine the relationship between the EEG signals and the behavioral pointing responses on a trial-to-trial level, we performed a representational similarity analysis between P-episode (which indicated oscillatory activity) for each event (Intersection 1, Leg 2, Leg 3, Intersection 2, Imagination) and the circular error of pointing responses. We calculated a 96 × 96 dissimilarity matrix between trials for each event period by using P-episode in the 1-30 Hz band and within our ROIs (Figure 2). For example, the P-episode in Trial (n+1) was correlated with that in Trial n by Pearson correlation. The correlation coefficient r was normalized as r_normalized_ = (1 - r)/2 and then were transformed to Fisher’s Z as Z = ln[(1 + r_normalized_) / (1 - r_normalized_)] / 2. We also calculated a 96 × 96 dissimilarity matrix between trials using the circular pointing error. The circular error in each trial was calculated as Error = Response direction – Predicted direction (also see Du et al., 2023). For each trial involving a pointing direction towards the participant’s left side (i.e., the correct pointing direction was in a range from 180° to 360°; which would involve trials with right-turn paths), we transformed the response pointing direction (ranging from 0° to 360°) to a range from −180° to 180° in which 0° was their facing direction at the end of the path. The sign on these values was then flipped to ensure these trials were comparable with the corresponding not-flipped trials for the same path (i.e., left-turn paths) and these values were normalized to range from 0° to 360°. The predicted pointing directions were also based on left-turn paths (i.e., the predicted pointing directions for Path 5-8 were in a range 0°-180°). Then, we normalized the circular pointing error to a range of [0°, 360°] and calculated the dissimilarity matrix of the circular pointing error between trials. For example, the dissimilarity was measured by the absolute value of the angular difference of the circular errors between Trial (n+1) and Trial n (normalized to [0°, 180°]).

**Figure 2.**
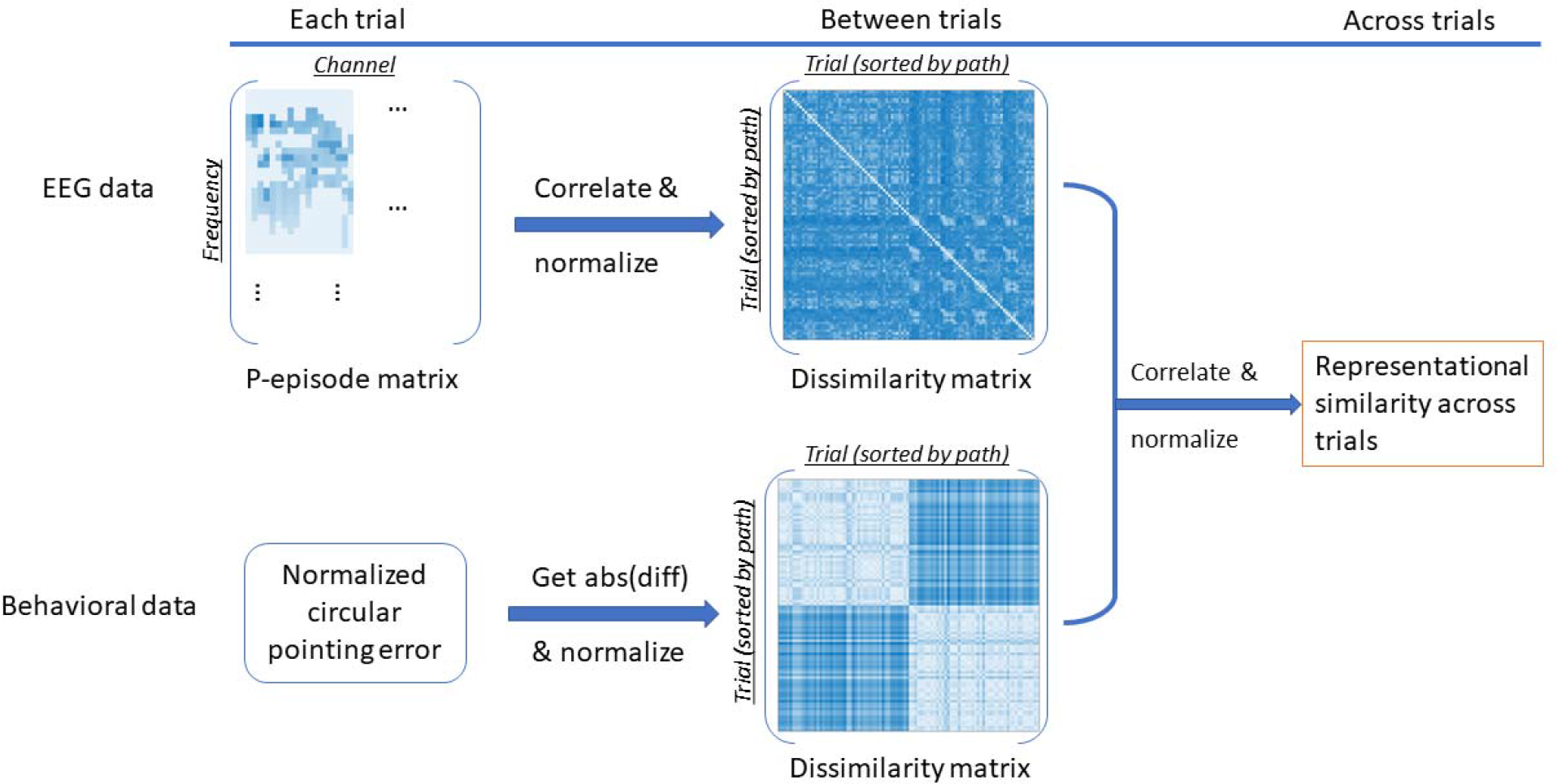
Calculation procedure for the representational similarity index between P-episode and circular pointing error in all trials.

We then performed partial correlations between the P-episode dissimilarity matrix and the pointing error dissimilarity matrix for each event (Intersection 1, Leg 2, Leg 3, Intersection 2, Imagination). For example, for the event Intersection 1, we performed a partial correlation between the dissimilarity matrix of Intersection 1 and that of pointing error while controlling the dissimilarity matrices of the other 4 events: Leg 2, Leg 3, Intersection 2, and imagination. Then we transformed the correlation coefficients to Fisher’s Z scores.

We compared the Fisher’s Z scores against 0 using t-tests. We also performed planned comparisons between these event pairs: Intersection 1 vs Leg 2, Intersection 2 vs Leg 2, Intersection 1 vs Leg 3, and Intersection 2 vs Leg 3.

#### 2.5.6. Representational similarity analysis within and between paths

We examined the representational similarity within the same path and between different paths by correlating P-episode between trials for each event period (Intersection 1, Leg 2, Leg 3, Intersection 2, Imagination). For example, the P-episode (in 1-30 Hz and within our ROIs) in Trial (n+1) was correlated with that in Trial n using Pearson correlation. The correlation coefficients r were then transformed into Fisher’s Z scores. We averaged the normalized similarity index across trials for each path to get the similarity index for each possible path pair. We then separated and averaged across path pairs in three categories: the same path in the same condition (e.g., Path 1 vs. Path 1; both in TI-C condition), different paths in the same condition (e.g., Path 1 vs. Path 2; both in TI-C condition), and different paths in different conditions (e.g., Path 1 in TI-C condition vs. Path 5 in FI-NC condition). We compared the similarity indices in the three categories.

#### 2.5.7. Classification analyses

We used a binary classifier to decode the trial conditions (TI-C or FI-NC) with the P-episode during each event (Intersection 1, Leg 2, Leg 3, Intersection 2, Imagination) for each frequency band. A binary logistic classifier was trained using Python *Scikit Learn* package. The ratio of train–test split for each iteration was 70–30%. For features used for training the classifier, we used the P-episode averaged across the electrodes in our ROIs within each trial for each event period. The training–testing sampling procedure was reiterated 100 times for each participant. The classification performance was evaluated by predictive accuracy. The mean decoding accuracy scores across all participants were compared with the chance level by two-tailed Wilcoxon signed rank tests. The chance level of decoding accuracy for each event and frequency band was calculated by shuffling labels (i.e., randomly selecting half of the trials to be TI-C condition and the remaining ones to be FI-NC) and running the same procedure as above.

For all analyses, all tests were two-tailed and used α = .05. FDR correction was applied where applicable. No part of the study analyses was pre-registered prior to the research being conducted.

## 3. Results

The behavioral pointing responses were reported in a previous study (Du et al., 2023). For the purposes of composition, we present the example pointing responses in two paths and the group circular pointing errors in Figure 1F, G, H, I. No trials were excluded from calculating mean circular pointing errors except those missing due to HMD disconnection.

EEG data and analysis code are available at https://osf.io/3aygm/files/osfstorage.

### 3.1. Power and oscillatory differences between conditions during walking and imagination phases

We first examined the EEG power when participants walked through the hallway and when they imagined where the start was after walking. Topographic plots of theta and alpha z-scored P-episode in true intersection – cross (TI-C) and false intersection – no cross (FI-NC) conditions are shown in Figure 3A-D. Mean power during walking and imagination was computed by averaging the channels in our ROIs for each condition (Figure 3E-F).

**Figure 3.**
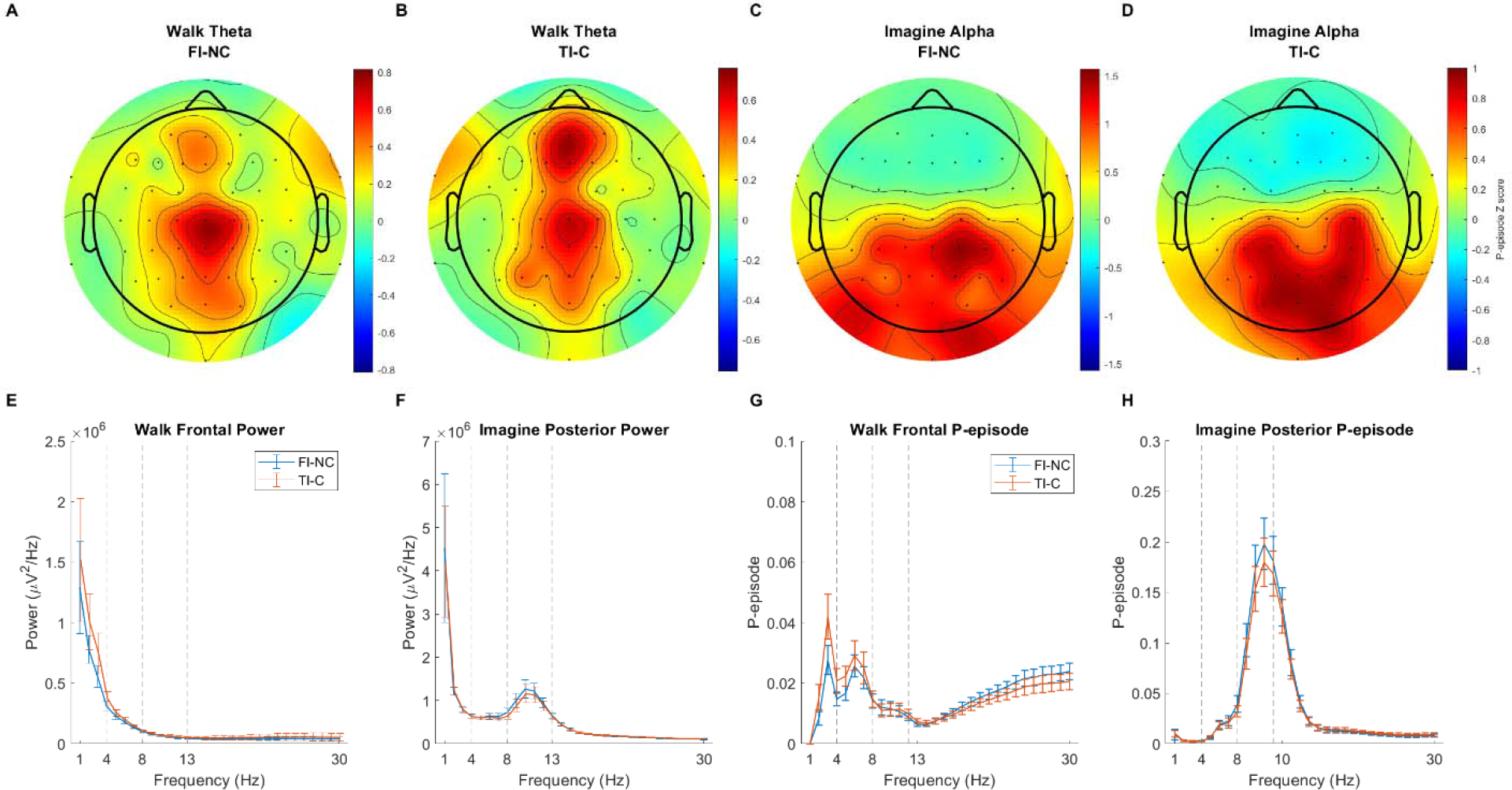
(A-D) Spatial distributions of theta and alpha oscillations (z-scored P-episode) in TI-C and FI-NC conditions. (E-F) Power spectral density as a function of frequency (Hz) in frontal and posterior ROIs. (G-H) P-episode as a function of frequency (Hz) in frontal and posterior ROIs. Blue lines show FI-NC condition. Red lines show TI-C condition. Error bars show the standard error of mean within the condition.

To investigate oscillatory activity, we computed mean P-episode across each event period for each ROI. For the walking phase, we then averaged across all event periods. The mean P-episode for each frequency band were shown in Figure 3G-H. For each frequency band and ROI (frontal theta and posterior alpha), a 2 (Event: walking/imagination) × 2 (Condition: TI-C/FI-NC) repeated-measure ANOVA was performed to compare the P-episode difference. Using the P-episode at theta band across the frontal regions, none of the main effects or the interaction was significant (all *p*s > .10, η_p_ ^2^ < .07). Planned comparisons showed that during walking, there was no significant difference between TI-C condition (Mean = 2.24 ×10^-2^, SD = 1.95 ×10^-2^) and FI-NC condition (Mean = 1.86 ×10^-2^, SD = 1.27 ×10^-2^), *t*(38) = 1.56, *p* = .127, Cohen’s *d* = 0.35. During imagination, there was no significant difference between the conditions, *t*(38) = 0.03, *p* = .980, Cohen’s *d* < 0.01 (FI-NC: Mean = 1.97 ×10^-2^, SD = 1.77 ×10^-2^; TI-C: Mean = 1.97 ×10^-2^, SD = 1.86 ×10^-2^). These findings suggest that the mean P-episode did not vary as a function of condition across the entire path, suggesting that any differences that might emerge during early compared to later portions of the hallway segment would not be due to larger differences in oscillatory activity across the conditions.

We then investigated whether theta and alpha oscillations varied during the different phases of the task, e.g., walking compared to imagination. Using the P-episode at alpha band across the posterior regions, there was a significant main effect of event: P-episode during imagination phase was significantly higher than during walking, *F*(1, 38) = 47.03, *p* < .001, *η*_p_^2^ = .55. The main effect of condition was significant: P-episode in FI-NC condition was significantly higher than in TI-C condition, *F*(1, 38) = 5.54, *p* = .024, *n*_p_^2^. = .13. The interaction was not significant, *F*(1, 38) = 3.80, *p* = .059, *n*_p_^2^. = .09. Planned comparisons showed that during imagination phase, the P-episode in FI-NC condition (Mean = 16.39 ×10^-2^, SD = 13.18 ×10^-2^) was significantly higher than that in TI-C condition (Mean = 14.77 ×10^-2^, SD = 12.54 ×10^-2^), *t*(38) = 2.23, *p* = .031, Cohen’s *d* = 0.51. During walking, there was no significant difference between the conditions, *t*(38) = 1.51, *p* = .139, Cohen’s *d* = 0.34 (FI-NC: Mean = 2.74 ×10^-2^, SD = 2.61 ×10^-2^; TI-C: Mean = 2.45 ×10^-2^, SD = 2.10 ×10^-2^). These results suggest that alpha oscillations were increased during the imagination compared to the walking period, consistent with our earlier findings on posterior alpha and frontal midline theta oscillations recorded with wireless scalp EEG (Liang et al., 2018). In addition, our results suggest that P-episode differed between the FINC and TI-C conditions during the imagination phase but not for the entire walking phase. These differences suggest that condition-specific information might be resolved toward the end of navigation, although looking at the entire walking period was likely too coarse to identify any condition or path specific differences.

### 3.2. True intersection areas were represented more similarly than false intersections within each path

To examine how different periods of the walking phase were represented, we used the P-episode (which indicated oscillatory activity) in individual trials to compute a representational similarity index between the walking events (see Section 2.5.4). We then averaged the similarity index (i.e., the Fisher’s Z scores) across trials in each condition (Figure 4). Representational similarity allowed us to compare neural similarity between different events in the same condition to determine how participants might be using the individual segments to represent the global path they walked on that trial. Specifically, we examined whether the neural representations between the events of the first and the second intersections (which looked identical) were different between the conditions. For both frontal and posterior regions, the similarity between the events of Intersection 1 and Intersection 2 in true intersection – cross (TI-C) condition was significantly higher than that in false intersection – no cross (FI-NC) condition (frontal: *t*(38) = 2.21, *p* = .043 FDR corrected, Cohen’s *d* = 0.50 (medium effect); posterior: *t*(38) = 2.50, *p* = .034 FDR corrected, Cohen’s *d* = 0.57). We also examined whether the representations between the events of the middle path segments (Leg 2 and Leg 3; which also looked identical) were different between the conditions. For frontal regions, the similarity level between the events of Leg 2 and Leg 3 in TI-C condition was significantly higher than that in FI-NC condition, *t*(38) = 2.09, *p* = .043 FDR corrected, Cohen’s *d* = 0.47. For posterior regions, there was no such effect, *t*(38) = 0.27, *p* = .791 FDR corrected, Cohen’s *d* = 0.06.

**Figure 4.**
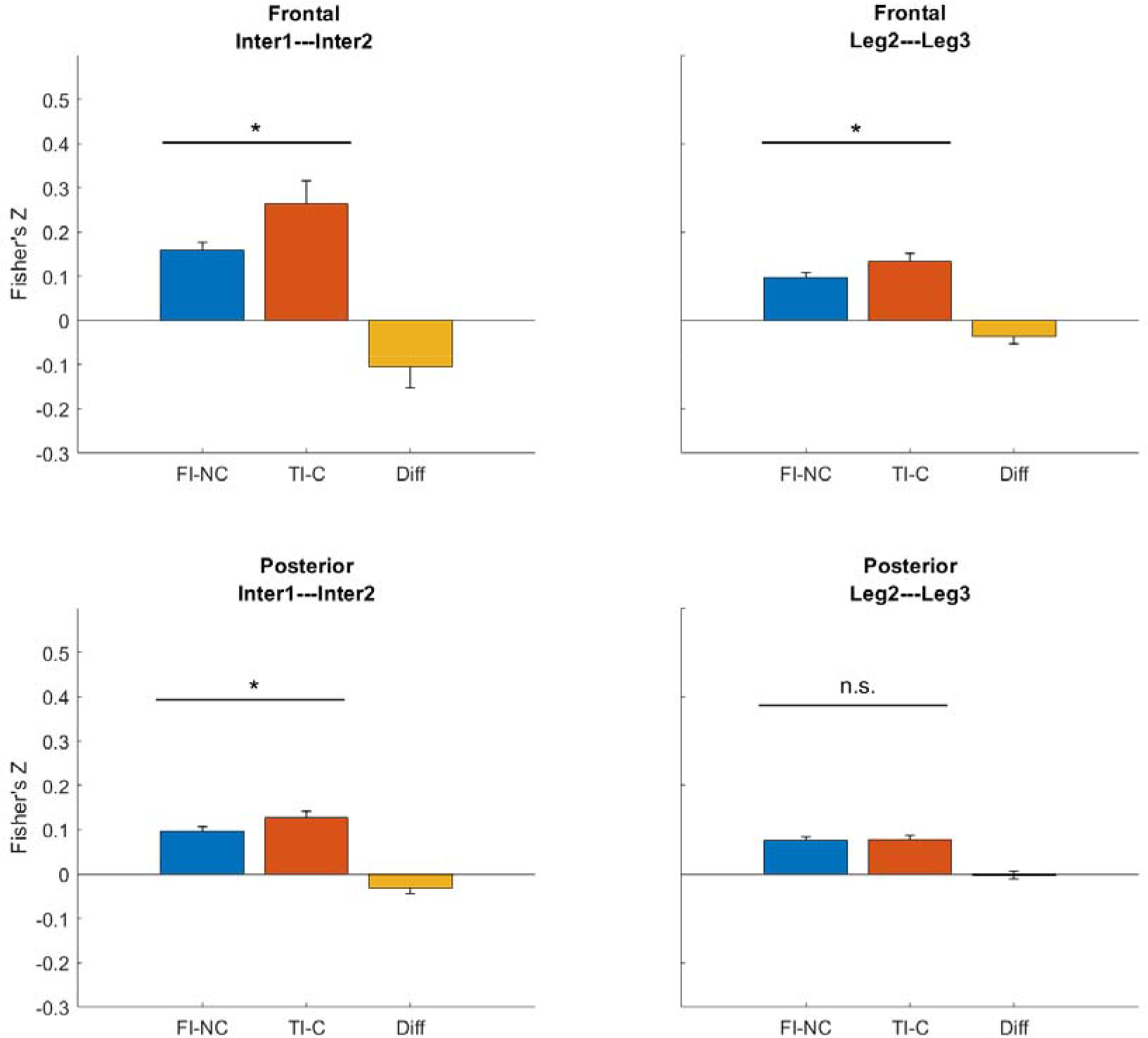
Representation similarity between events using P-episode. Blue bars show FI-NC condition. Red bars show TI-C condition. Yellow bars show the difference between conditions (FINC-TIC). Error bars show the standard error of mean within the condition. *: *p* < .05. n.s.: *p* > .05 (FDR corrected).

These results suggest that in the TI-C condition, in which there was no conflict when walking through the first and the second intersections, the neural representations for the identical-looking segments were more similar than in the FI-NC condition. This information would be important to participants’ subsequent decisions because in the TI-C condition, there was no violation of physically crossed paths: Leg 1 and Leg 4 actually crossed (Figure 1). In contrast, in the FI-NC condition, Leg 1 and 4 never actually crossed despite the presence of two false intersections giving the illusion that they crossed. Notably, neural representations were also more similar during the middle path segments in the TI-C condition than in FI-NC condition. Based on these findings, we next examined whether visual information experienced early during navigation – as early as Leg 1 (when the first intersection was presented) contributed to subsequent pointing patterns.

### 3.3. Oscillatory activity during walking and imagination positively correlates with circular pointing error on each trial

The evidence obtained so far from wireless scalp EEG recordings suggests that participants may have been reconciling competing information during the imagination phase but that representations related to the paths taken emerged as early as the first leg when the intersection was visible. We next examined how representational similarity related to subsequent behavioral pointing response, wherein participants gave their responses about the path they walked. Pointing error also provided an index of whether participants created a crossed path from an uncrossed path in the FI-NC condition (Du et al., 2023). To examine the relationship on a trial-to-trial level (i.e., both TI-C and FI-NC trials were included), we computed a representational similarity index between P-episode and the circular error of pointing response by performing partial correlations between the matrices (Figure 2; see Section 2.5.5). For both frontal and posterior regions, the similarity levels in all events were above 0 (*t*s > 3.14, *p*s < .004 after FDR correction, Cohen’s *d*s > 0.71), suggesting a positive association between oscillatory activity during encoding and the following memory retrieval accuracy.

To gain insight into when neural representations related to the shapes of paths they walked first emerged, we examined whether walking at the intersection areas was more strongly associated with the retrieved memories than in the middle path segments. Using the calculated similarity indices between P-episode at each epoch and the subsequent pointing error at the end of the hallway, we compared between different event periods: Intersection 1 vs. Leg 2, Intersection 1 vs. Leg 3, Intersection 2 vs. Leg 2, and Intersection 2 vs. Leg 3. For frontal regions, planned comparisons showed that the similarity levels in the intersection areas were significantly higher than that in the Leg 2 or Leg 3 events: Intersection 1 vs. Leg 2, *t*(38) = 2.84, *p* = .028, Cohen’s *d* = 0.64; Intersection 1 vs Leg 3, *t*(38) = 2.27, *p* = .038, Cohen’s *d* = 0.52; Intersection 2 vs Leg 2, *t*(38) = 2.32, *p* = .038, Cohen’s *d* = 0.52; Intersection 2 vs Leg 3, *t*(38) = 1.64, *p* = .110, Cohen’s *d* = 0.37 (all *p*s FDR corrected). For posterior regions, none of these comparisons reached significant difference (*t*s < 2.02, *p*s > .204 after FDR correction, Cohen’s *d*s < 0.46). These results indicate that the frontal-midline oscillatory activities when walking through the first or the second intersections were more correlated with the retrieved memories than when walking in the middle segments of the path. These findings suggest that neural representations correlate (albeit weakly) with subsequent pointing patterns and that such neural representations emerge early at intersections rather than when simply walking.

We also performed the same analysis using each of the 1-second periods of the imagination phase separately (see supplementary materials).

### 3.4. Paths within the same condition are represented more similarly than paths in different conditions

The representational similarity analysis above (Section 3.3) showed that across different paths and conditions, the oscillatory activities during encoding were positively related to the following memory error. Although we had two different conditions (TI-C and FI-NC), we employed a total of four different paths in each condition. While the conditions were blocked, the paths themselves were randomized within blocks such that participants would not be able to guess their global path (and therefore their end location) as they walked. One may argue that all paths were quite similar to each other so that, regardless of conditions, they were always represented similarly and were always correlated with the memory error. Although the paths were clearly of different shapes (see Du et al., 2023, Table S1 and Figure S1 for details), to test this possibility at the neural level, we performed a similarity analysis by correlating P-episode between trials and averaging the normalized similarity index across path pairs. We did this for three different categories^1^: the same path in the same condition (e.g., Path 1 vs. Path 1; both in TI-C condition), different paths in the same condition (e.g., Path 1 vs. Path 2; both in TI-C condition), and different paths in different conditions (e.g., Path 1 in TI-C condition vs. Path 5 in FI-NC condition; see Section 2.5.6). If all paths were represented similarly regardless of paths or conditions, the similarity level should be equivalent in all categories. Otherwise, there should be significant differences between the categories.

Figure 6 shows the mean similarity indices of the three categories. The similarity level for the same path category (blue dotted lines) was significantly higher than the other two categories in all event periods for both frontal and posterior regions, *t*s > 22.56, *p*s < .001 (FDR corrected), Cohen’s *d*s > 5.12. For frontal regions, the similarity level for the different paths in the same condition (red dashed lines) was significantly higher than that for different paths in different conditions (green solid lines) in all event periods, *t*s > 5.21, *p*s < .001 (FDR corrected), Cohen’s *d*s > 1.18. For posterior regions, the similar effects were found in all event periods, *t*s > 3.63, *p*s < .001 (FDR corrected), Cohen’s *d*s > 0.82.

After separating the same path pairs, we found that the paths within the same condition were represented more similarly than the paths in different conditions, which suggests that the manipulation of path type was consistent with our expectations. This result rules out the possibility that the paths in different conditions were always represented similarly and therefore suggests that the positive relationship between neural oscillatory patterns and pointing errors in previous analysis (Section 3.3) was more likely to indicate an encoding-retrieval mechanism. We will revisit this point later in the discussion.

### 3.5. Successful decoding of trial condition from frontal-midline theta oscillations

To further examine the oscillatory difference between conditions, we trained and tested a binary classifier to decode trial conditions (TI-C vs. FI-NC) for each event (Figure 7 and S6; see Section 2.5.7). Using the P-episode at theta band in frontal regions, we found that the decoding accuracies in all events (including walking and imagination phases) were significantly higher than the chance level (Wilcoxon signed rank tests: *Z*s > 3.22, *p*s < .002, FDR corrected). Using the P-episode at alpha band in posterior regions, the decoding accuracies during walking in Intersection 1, Leg 3, and imagination phase were significantly higher than the chance level (Wilcoxon signed rank tests: *Z*s > 2.60, *p*s < .02, FDR corrected) whereas the decoding accuracies during walking in Leg 2 and Intersection 2 were not different from the chance level (Wilcoxon signed rank tests: *Z*s < 1.86, *p*s > .07, FDR corrected). These results indicate that differences in P-episode in frontal-midline regions were sufficient to classify condition during some events, even in the earlier stages of navigation (e.g., when walking through the area of the first intersection). These findings again support the idea that neural representations for the global path shape, as measured using oscillatory prevalence, emerged early during navigation.

## 4. Discussion

The current study examined how humans develop spatial memories for the global path that they walk when there is conflicting allothetic and idiothetic information. Using mobile scalp EEG recordings, we analyzed the oscillatory activities during navigation learning to test the serial vs. parallel learning hypotheses. We will discuss three different important aspects of our findings.

First, by correlating the P-episode with the pointing error, we found that encoding the intersection areas of the path was more strongly associated with the subsequent memory error than the middle legs (Figure 5). This effect was not simply due to similar path structures. In alignment with this idea, there was a prominent difference in representations for the intersection areas between the true intersection – cross (TI-C) and false intersection – no cross (FI-NC) conditions, but not in representations for the middle path segments (Figure 4). This suggests a more salient role for the intersection areas in contributing to the memory of the path.

**Figure 5.**
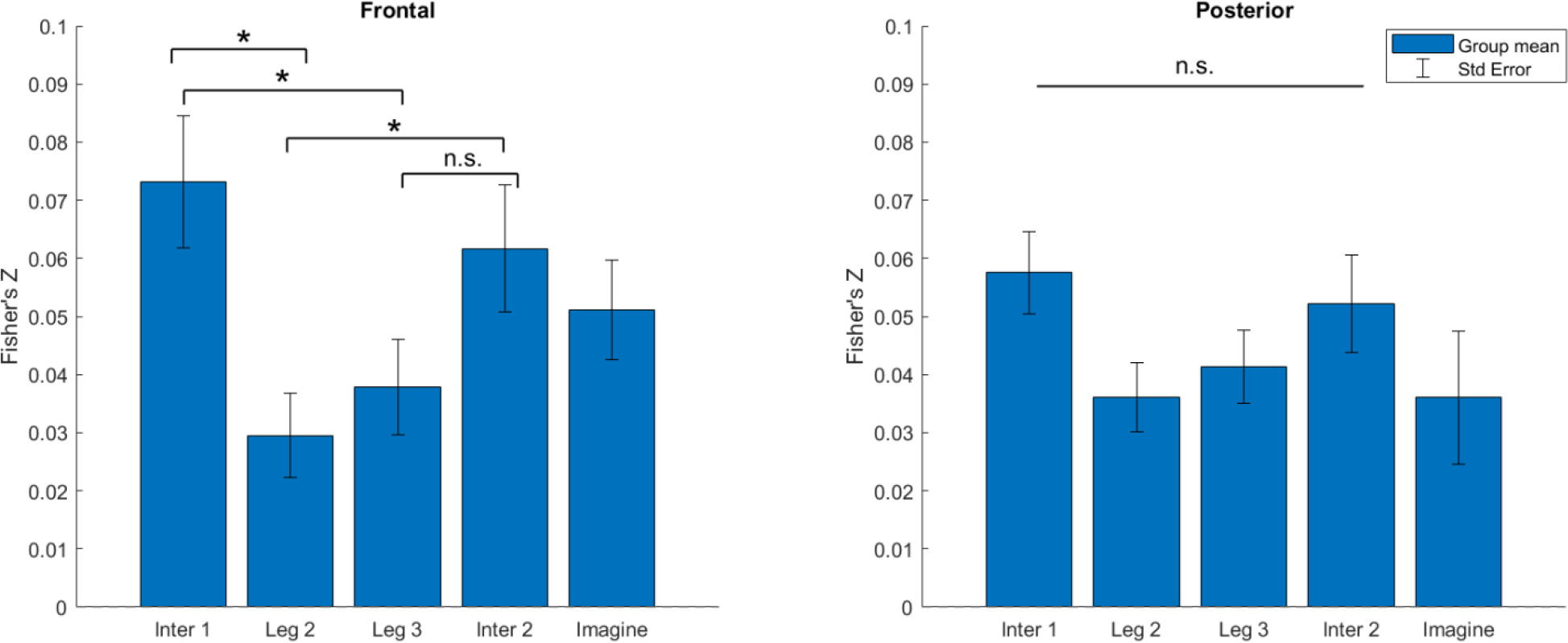
Representational similarity between P-episode and circular pointing error in all trials using frontal and posterior P-episode. Error bars show the standard error of mean within the condition. *: *p* < .05. n.s.: *p* > .05 (FDR corrected).

Furthermore, the mean classification decoding accuracies in earlier stages of navigation (Intersection 1 and Leg 2) were significantly above chance level (Figure 7), which suggests that the participants started to make inferences about the path shape at a quite early stage of navigation. These results are consistent with our parallel learning hypothesis rather than the serial learning hypothesis. Encoding early or late throughout the navigation does not determine the importance or the reliability of any navigation portions. Our results clearly support that the intersection areas, even the first intersection in the first leg of the path, showed a relatively greater weighting than the middle segments in forming the memory of the path. The intersections were salient features along the hallway so that they might have served as memory anchors and weighed more than other segments in the learning process.

Second, the current study included a conflict condition (FI-NC) and a non-conflict condition (TI-C) to test when the combination of allothetic and idiothetic information occurred. By comparing the P-episode during imagination, we found significantly greater alpha oscillations in FI-NC condition than in TI-C condition, which could relate to the greater need for reconciling memories for the conflicting information acquired during navigation in the false intersection – no cross (FI-NC) condition compared to the true intersection – cross (TI-C) condition. Notably, the oscillatory patterns in the first and the second intersection areas were more similar to each other for the TI-C condition than the FI-NC condition (Figure 4), which suggests the information in both intersection areas was involved in the computations of path shape. The representational similarity analysis within and between path types further supports this idea that starting from the early stages of navigation, there was a clear difference in the oscillatory patterns between the paths in the same condition than those in different conditions (Figure 6). This suggests that the representation of the path involved the information acquired at both the first intersection and later portions of navigation including the second intersection. Overall, given that the oscillatory patterns we recorded showed a difference early on during navigation, and likely related to memory and other decision-making processes that occur throughout navigation, our findings are more consistent parallel hypothesis.

**Figure 6.**
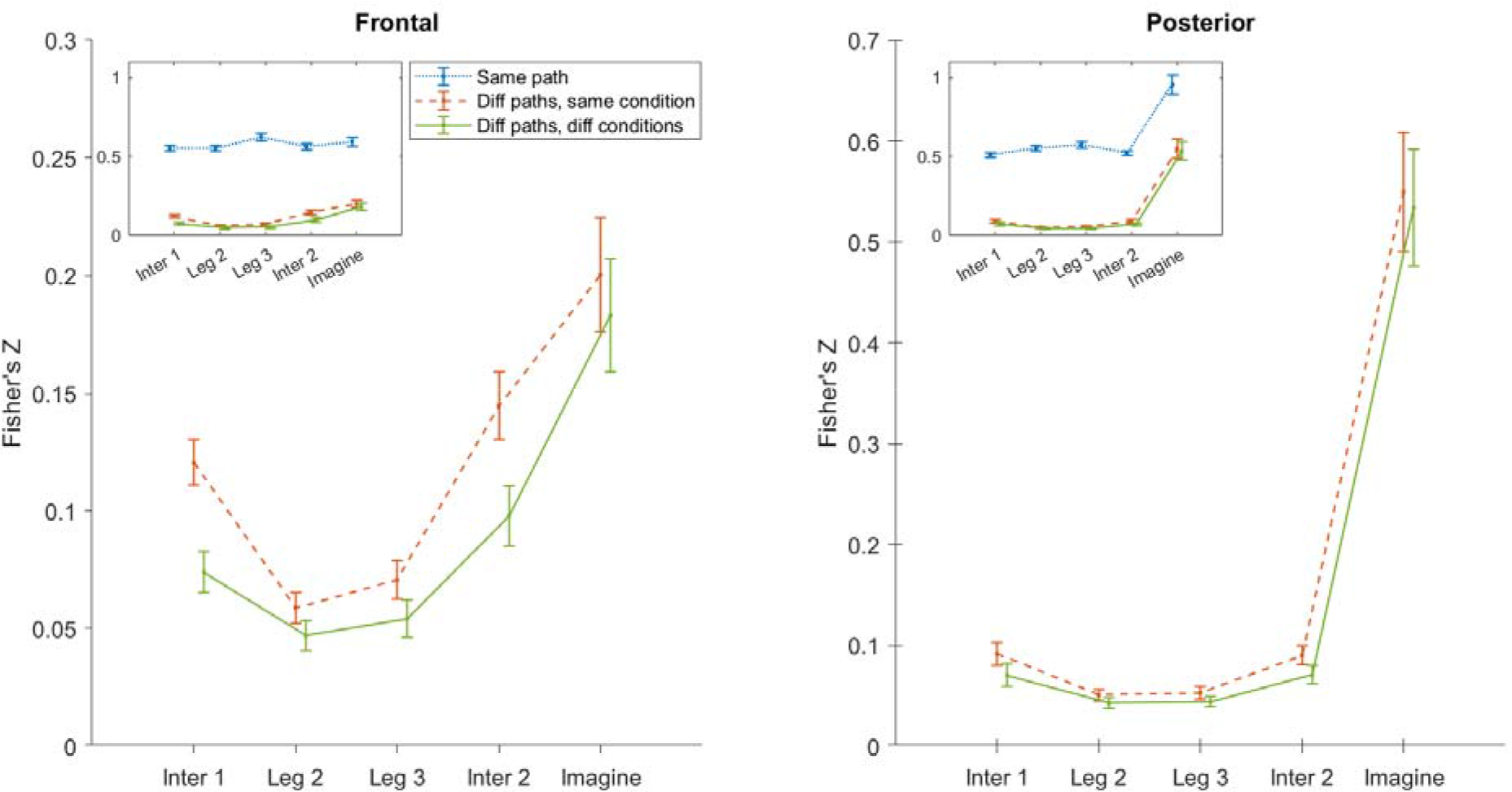
Representational similarity within the same path, the same condition, and between conditions using frontal and posterior P-episode. Blue dotted lines show the mean of path pairs for the same path. Red dashed lines show the mean of path pairs for different paths in the same condition. Green solid lines show the mean of path pairs for different paths in different conditions. Error bars show the standard error of mean within the condition.

We speculate that in the early stages of navigation, participants may have formed multiple representations and predictions about the path based on their individual and past-trial experience, extending our previous proposal based on behavioral results (Du et al., 2023). For example, during the navigation in the first intersection area, participants might try to predict the path shape according to their visual or body-based input. These representations are not always consistent (i.e., in conflict, like in the FI-NC condition) and may compete or combine with each other across the navigation learning phase until memory retrieval is required (the pointing phase) (L. Zhang & Mou, 2017). Our findings are consistent with previous studies proposing an active process of comparing available information with current predictions (Harootonian et al., 2020; Ishikawa & Montello, 2006; Nardini et al., 2008; Sjolund et al., 2018; H. Zhang et al., 2014; Zhao & Warren, 2015). When there are multiple sources of information, there is a need to compare memories and make decision about idiothetic and landmark information, much of which likely occurs without the navigator’s explicit knowledge. We hypotheses that such cognitive processes likely involve different weightings of the sources (e.g., different cues in the environment, different modalities of input, etc.), possibly following Bayesian probabilistic models (Nardini et al., 2008; Newman et al., 2023; L. Zhang et al., 2020).

Third, we also examined the role of theta (4-8 Hz) oscillations in frontal-midline regions in the current study. Consistent with previous research (Chrastil et al., 2022; Liang et al., 2018, 2021), we found that theta oscillatory activity during active movements (the walking phase) was more salient in frontal-midline regions than in other regions. Frontal-midline theta oscillations during walking did not differ between conditions, consistent with our previous results suggesting that movement is one driver of frontal-midline theta oscillations (Liang et al., 2018). Our findings also suggest that frontal-midline theta oscillations were associated with encoding spatial information in active navigation and not simply movement as they contained information about condition and path shape. This is because frontal-midline theta oscillations were more correlated in the TI-C compared to FI-NC condition and were sufficient for classifying one condition compared to another, even as early as the first intersection.

We also observed that alpha oscillations during the imagination phase (prior to the pointing response) were more salient in parietal and occipital regions than other regions. Since the participants in the imagination phase were asked to compute the start’s position with the memory of the walked path to prepare for the following motoric actions (pointing), the increased alpha oscillations may suggest increased attention prior to decision making as in previous research with virtual navigation tasks (Chrastil et al., 2022). Another possibility is that alpha oscillations reflected inhibition of competing information (Waldhauser et al., 2012) such as conflicting idiothetic cues, which would be expected to be greater during FI-NC than TI-C paths. In the classification analysis, comparing to the observed posterior alpha oscillations, the frontal-midline theta oscillations were better indicators to distinguish between the true intersection – cross (TI-C) and false intersection – no cross (FI-NC) conditions across the active navigation periods and the imagination phase (Figure 7). These results further support a mnemonic role of theta frontal-midline oscillations in forming spatial memories in previous research (Cornwell et al., 2008; Ekstrom & Watrous, 2014; Kaplan et al., 2012).

**Figure 7.**
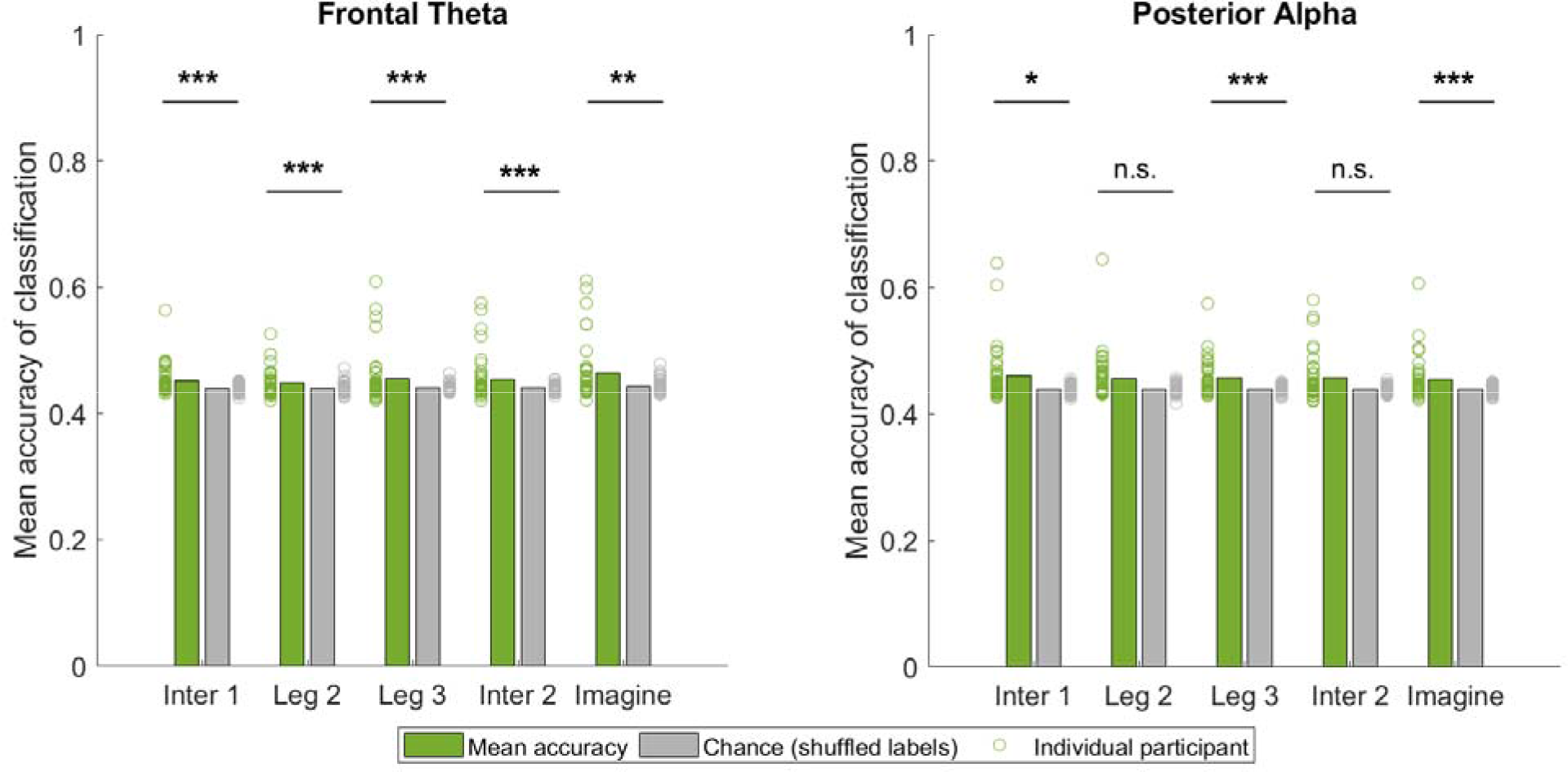
Classification accuracy by using frontal theta and posterior alpha P-episode. Bars show the group mean across participants. Dots show the accuracy of individual participants. ***: *p* < .001. **: *p* < .01. *: *p* < .05. n.s.: *p* > .05 (FDR corrected).

One may argue that it is not surprising to see the first intersection portion was related with later memory because trials in the same condition were presented in blocks. That is, participants learned “this is a TI-C/FI-NC path” from previous trials rather than forming representations solely on the current path (although they were never told that there were two conditions). However, while the blocks may have provided some ability to better extract condition specific regularities, the early-parallel-learning effects were unlikely to be influenced solely by the blocked design. It is important to note that the path shapes themselves were randomized within blocks (see methods) so participants would be unable to guess the global path shape or their end location. Also, the first path segment varied in length across paths in both conditions (see Table S1 and Figure S1) such that participants would not know the global path shape when they began navigating that trial. The participants may have made guesses according to previous trials in the same condition, but representing the current path shape accurately would require encoding of the specific path. The length of L1 alone did not provide sufficient information to estimate the pointing angle at the end of the navigation (see Figure S1). We also note that our previous behavioral results were consistent using a blocked design and random/mixed design in different groups of participants (Du et al., 2023). Representational similarity analyses (Figure 6) showed that the same paths were more similar -- on neural representations -- than different paths, suggesting that path shape contributed to the emerging representations as they navigated.

While length was unlikely to be the sole factor guiding participants’ early decisions about their start location, this begs the question what factors may have driven differences in FI-NC and TI-C conditions even by the first leg. While this was not explicitly manipulated in our design, we speculate that participants may have extracted regularities about intersection position and other features of the path shapes over the block that could have informed their (likely implicit) decision. Future experiments could address what factors might drive participant navigational decisions early during learning although we think that this study makes clear that participants were likely making such decisions earlier than might be expected based on (classic) serial models. Another potential limitation that could be addressed in future experiments is that scalp EEG may provide only limited insight into the neural basis of relevant cognitively relevant decision-making during navigation. This limitation is more difficult to directly address because mobile experiments such as performed here are not possible with other methods such as fMRI. Implanted wireless recordings may offer one possible solution although the coverage of these is somewhat limited (Stangl et al., 2023).

Together, our observations using wireless scalp EEG during active navigation indicate representations of a path likely form during early stages of navigation rather than during late stages alone, although clearly, some important aspects of decision making for conditions happen during late periods as well. Our results suggest that frontal-midline theta oscillations during active navigation directly, in particular, relate to such decision and memory making processes, possibly indicating the mnemonic role of theta oscillations.

## Supporting information

All supplements

## Acknowledgements

We thank the following research assistants for their contribution to this study: Kayla Cao, Anna Ramsook, Cameron Dockens, and Stephanie Doner. We thank Dr. Li Zheng for useful discussion and comments on the data analysis.

Funding information: NSF BCS-1630296 to ADE, NIH/NINDS R01NS076856 to ADE.

## Footnotes

1 There was no combination of “same path in different conditions” in this study. This was because there were four path types in each condition (TI-C vs FI-NC) and all of them were necessarily different from each other (see Figure S1 and Table S1). Path number was embedded in condition.

