## Supplementary material for "Frontal-midline theta and posterior alpha oscillations index early processing of spatial representations during active navigation": All supplements

**Supplementary Materials**

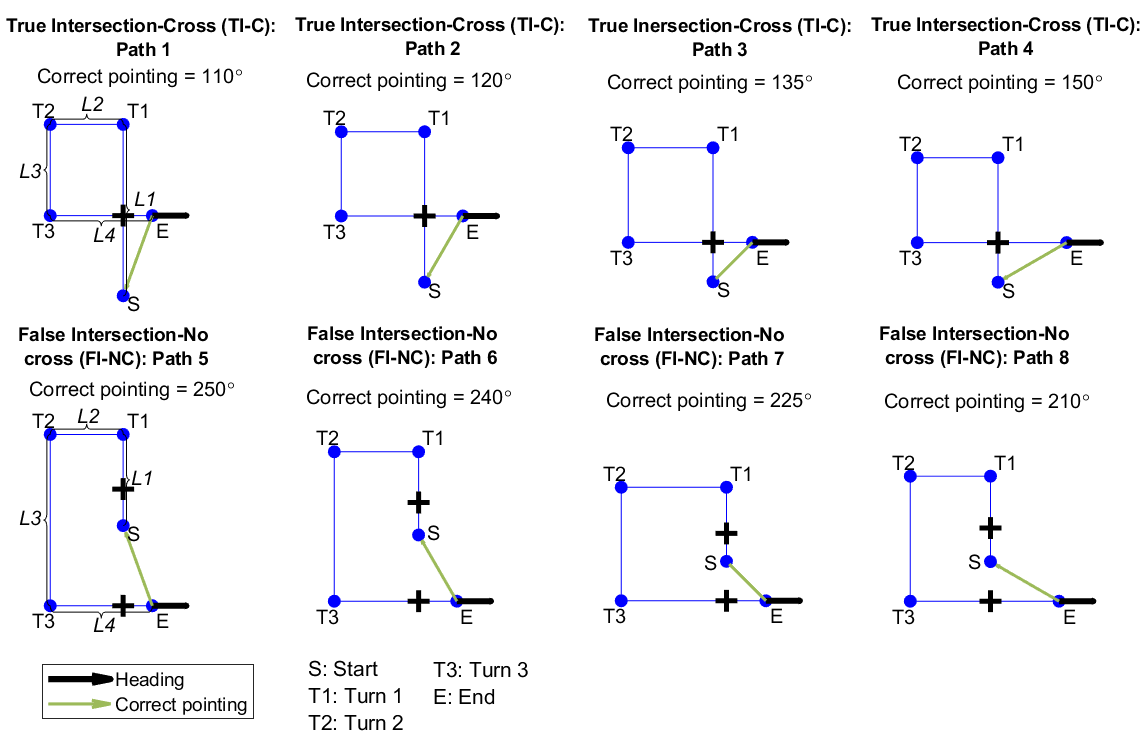

*Figure S1*. Path shapes and predicted correct pointing directions. Green arrows indicate the correct pointing direction according to Euclidean geometry. Bold pluses indicate the position of the intersections. S: Start. T1-3: Turn 1-3. E: End.

Table S1. *Path dimensions (in meters).*

| **Path** | **Path type** | **L1** | **L2** | **L3** | **L4** | **Total length** |
| --- | --- | --- | --- | --- | --- | --- |
| 1 | Cross | 3.99 | 1.70 | 2.12 | 2.38 | 10.2 |
| 2 | Cross | 3.49 | 1.93 | 1.95 | 2.82 | 10.2 |
| 3 | Cross | 3.12 | 1.98 | 2.20 | 2.90 | 10.2 |
| 4 | Cross | 2.89 | 1.87 | 1.97 | 3.47 | 10.2 |
| 5 | No cross | 2.12 | 1.70 | 3.99 | 2.38 | 10.2 |
| 6 | No cross | 1.93 | 1.96 | 3.47 | 2.84 | 10.2 |
| 7 | No cross | 1.73 | 2.45 | 2.65 | 3.37 | 10.2 |
| 8 | No cross | 1.98 | 1.86 | 2.90 | 3.45 | 10.2 |

*Note*: L1: From the start (S) to the first turning point (T1). L2: From T1 to the second turning point (T2). L3: From T2 to the third turning point (T3). L4: From T3 to the end (E). See Figure S1 for more details.

**S1. Mean power and P-episode at alpha band from posterior ROIs for the imagination phase**

We separated the whole imagination period (4 seconds) into four 1-second periods and calculated the mean power and P-episode within each period (Figure S2 and S3).

For the mean P-episode in alpha band within each period, 2 (Time: t1, t2, t3, t4) × 2 (Condition: FI-NC, TI-C) repeated-measure ANOVA showed that there was a significant difference across time, *F*(1.60, 60.81) = 3.93, *p* = .033 (Greenhouse-Geisser corrected), *η*_p_^2^ = .09. The main effect of condition was significant: P-episode in FI-NC condition was significantly higher than in TI-C condition, *F*(1, 38) = 4.38, *p* = .043, *η*_p_^2^ = .10. The interaction was not significant, *F*(2.91, 110.66) = 0.99, *p* = .397 (Greenhouse-Geisser corrected), *η*_p_^2^ = .03. Further analysis indicated that the mean P-episode in the second period was significantly higher than all other period, *t* > 2.71, *p* < .03 (FDR corrected), Cohen’s *d* > 0.61.

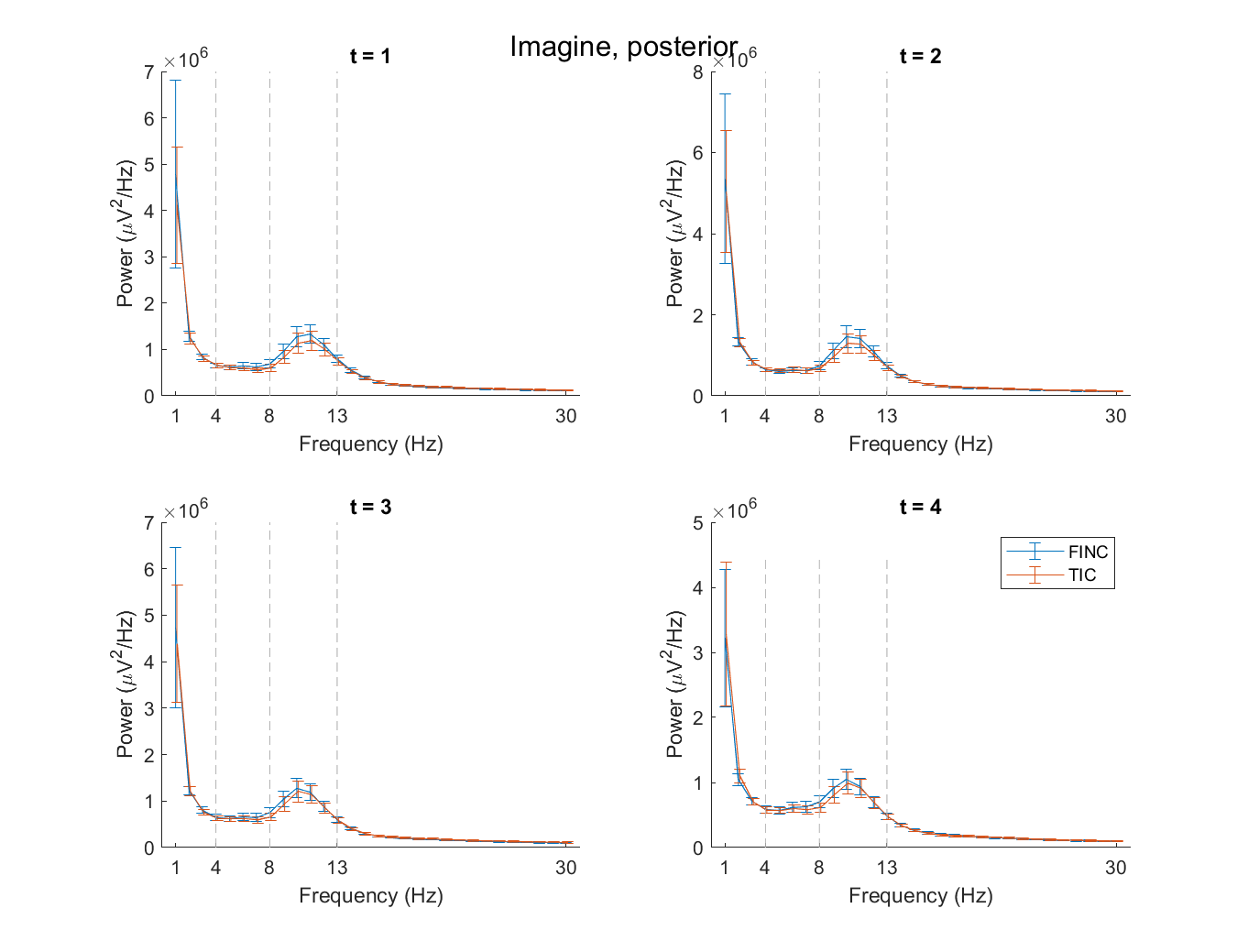

*Figure S2*. Power spectral density as a function of frequency (Hz) in posterior ROIs. No baseline correction was applied. t = 1: the first one second. t = 2: the second one second. t = 3: the third one second. t = 4: the last one second. Blue lines show FI-NC condition. Red lines show TI-C condition. Error bars show the standard error of mean within the condition.

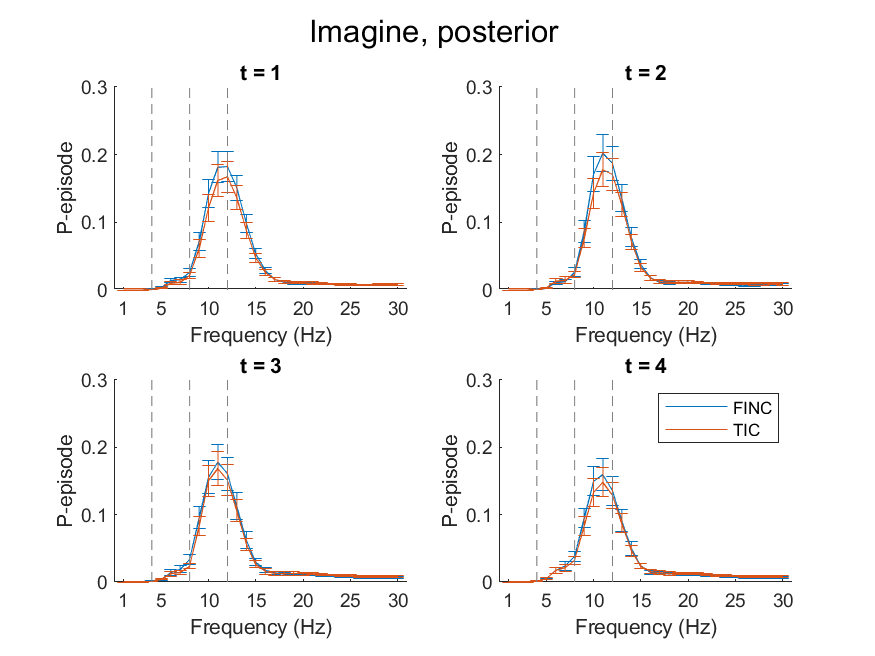

*Figure S3*. P-episode as a function of frequency (Hz) in frontal and posterior ROIs. t = 1: the first one second. t = 2: the second one second. t = 3: the third one second. t = 4: the last one second. Blue lines show FI-NC condition. Red lines show TI-C condition. Error bars show the standard error of mean within the condition.

**S2. Representational similarity between the P-episode within 1-second periods of the imagination phase and subsequent pointing error**

We separated the imagination period (4 seconds) into four 1-second periods and performed the representational similarity analysis between the P-episode and the subsequent circular pointing error (Figure S4 and S5). Note that the Fisher’s Z values for the walking events were slightly different between the periods because they were calculated based on partial correlation coefficients. For frontal ROIs, we did not find any significant differences between the four periods (repeated-measure ANOVA: *F*(2.34, 88.93) = 1.57, *p* = .211, *η*_p_^2^ = .04, Greenhouse-Geisser corrected). For posterior ROIs, repeated-measure ANOVA showed that there was a significant difference between the four periods, *F*(3, 114) = 2.83, *p* = .042, *η*_p_^2^ = .07. Further analysis indicated that there was a significantly stronger correlation between P-episode and subsequent pointing error for the first period compared to that for the second period, *t*(38) = 3.04, *p* = .024 (FDR corrected), Cohen’s *d* = 0.69. This may suggest that participants started to make decisions in early periods of the imagination phase.

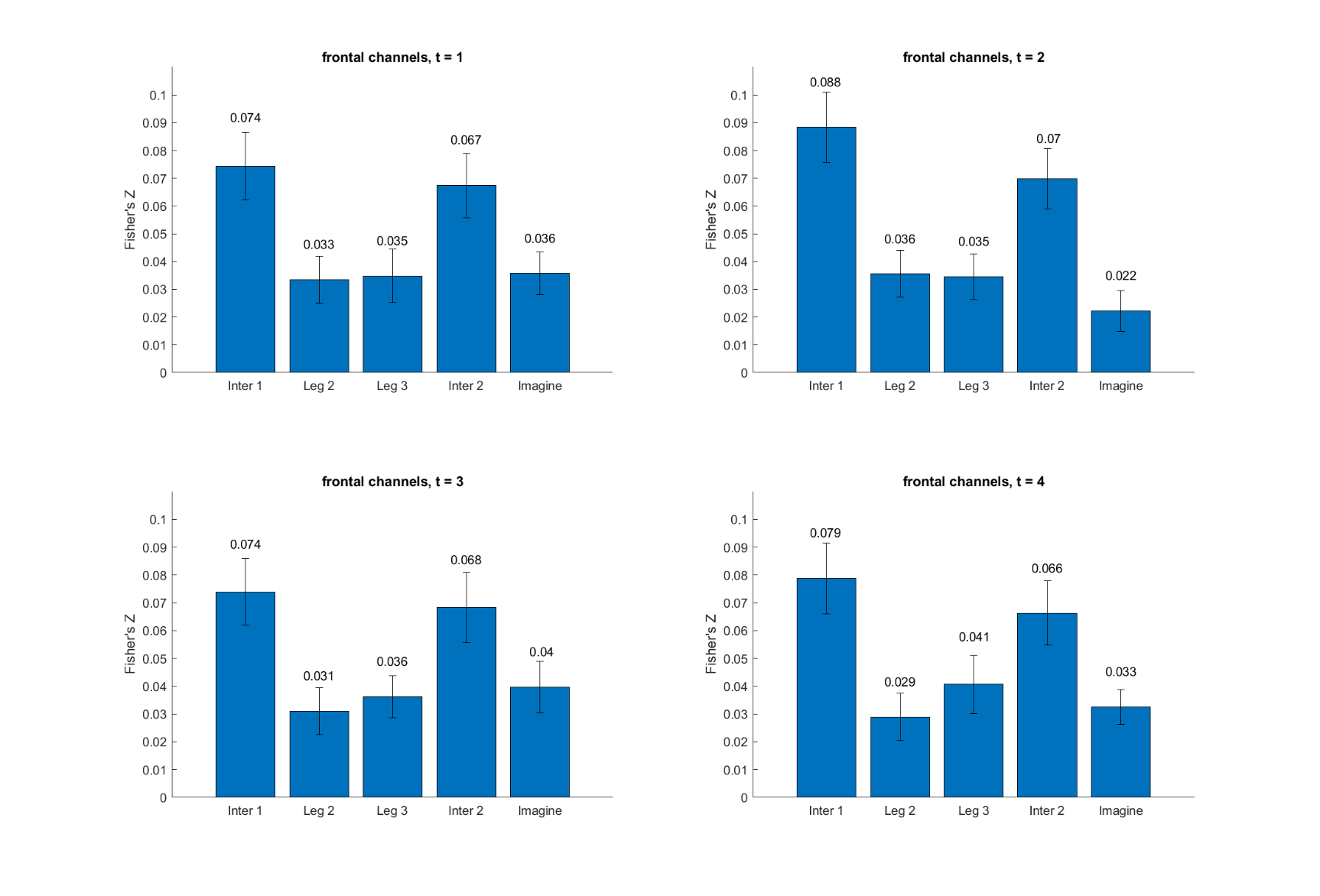

*Figure S4*. Representational similarity between P-episode and circular pointing error using 1-second periods of the imagination phase in frontal ROIs. t = 1: the first one second. t = 2: the second one second. t = 3: the third one second. t = 4: the last one second. Error bars show the standard error of mean within the condition.

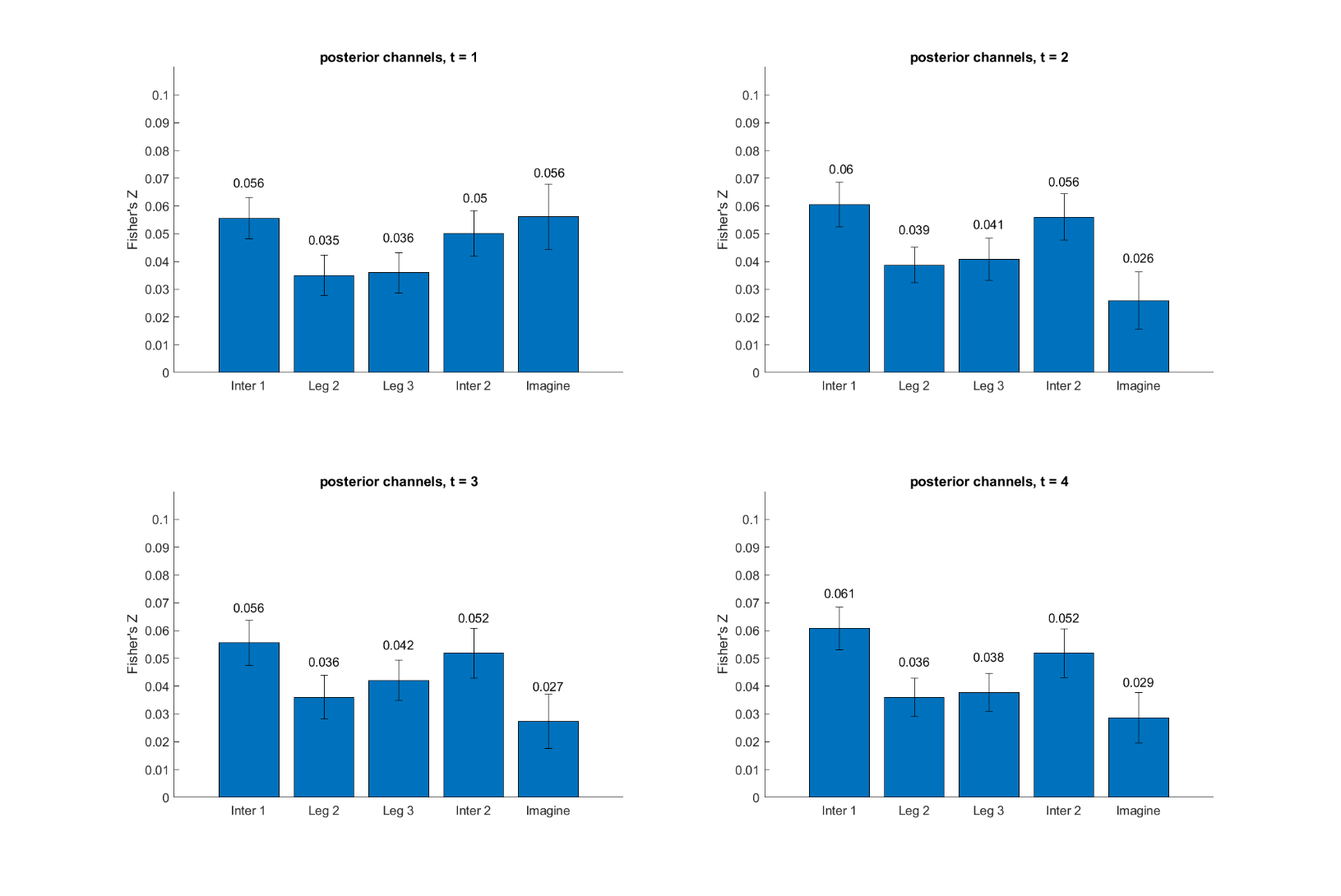

*Figure S5*. Representational similarity between P-episode and circular pointing error using 1-second periods of the imagination phase in posterior ROIs. t = 1: the first one second. t = 2: the second one second. t = 3: the third one second. t = 4: the last one second. Error bars show the standard error of mean within the condition.

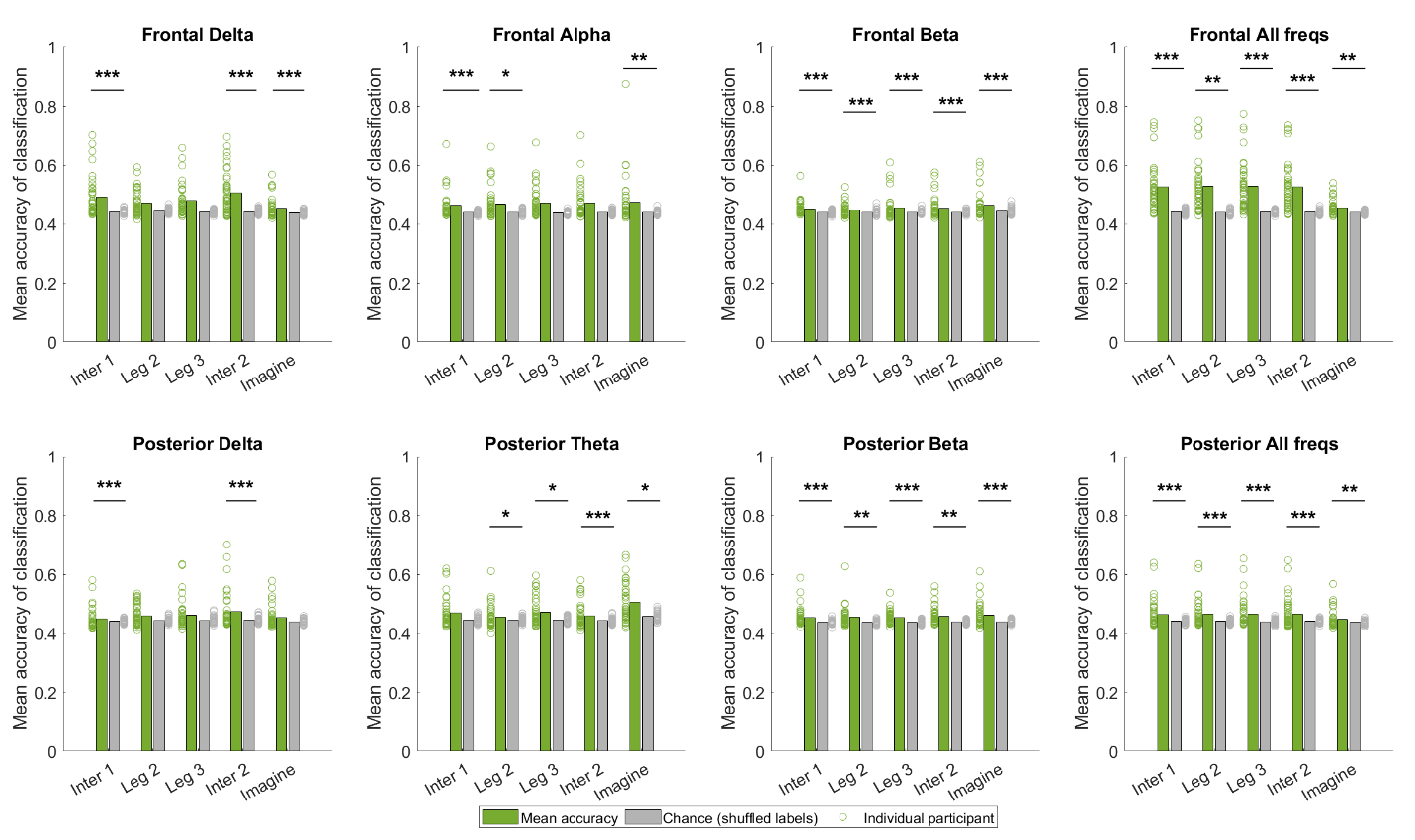

*Figure S6.* Classification accuracy by using frontal and posterior P-episode. Bars show the group mean across participants. Dots show the accuracy of individual participants. ***: *p* < .001. **: *p* < .01. *: *p* < .05 (FDR corrected).
